# Glucocorticoid Receptor Signaling in Myeloid Cells Orchestrates Inflammation Resolution and Muscle Repair

**DOI:** 10.1101/2025.10.29.685370

**Authors:** Sirine Souali-Crespo, Joe G. Rizk, Emilia Calvano, Rajesh Sahu, Emina Colovic, Imane Chabba, Valentine Gilbart, Erwan Grandgirard, Bastien Morlet, Qingshuang Cai, Daniel Metzger, Delphine Duteil

## Abstract

Glucocorticoids are key regulators of inflammation and tissue repair, yet their precise role in muscle regeneration remains incompletely understood. Here, we investigate the impact of myeloid-specific glucocorticoid receptor (GR) invalidation on macrophage dynamics and muscle stem cell function following acute injury. We demonstrate that the loss of GR in myeloid cells leads to increased macrophage accumulation, driven by altered proliferation and recruitment, without affecting fibro-adipogenic progenitor differentiation or satellite cell proliferation and differentiation under steady-state conditions. Transcriptomic and cistrome analyses at early regeneration stages reveal that GR directly regulates gene networks involved in efferocytosis and cell cycle control in myeloid cells. Importantly, administration of dexamethasone during the pro-inflammatory phase markedly delays muscle regeneration by impairing monocyte-to-macrophage transition and promoting macrophage proliferation in a myeloid-GR dependent manner, ultimately reducing satellite cell proliferation and myogenesis. In contrast, dexamethasone treatment during the anti-inflammatory phase exerts limited effects on muscle recovery. Together, our findings uncover a critical temporal role of GR signaling in myeloid cells in coordinating inflammatory resolution and stem cell function during muscle repair, and highlight the complexity of glucocorticoid actions in regenerative contexts.

## Introduction

Skeletal muscle injury is a common clinical challenge with profound effects on mobility and life quality. Muscle regeneration is orchestrated through a tightly regulated interplay between tissue-resident muscle stem cells, known as satellite cells (SCs) and their surrounding niche, which includes fibroadipogenic progenitors (FAPs), fibroblasts, endothelial cells, and immune cells^1–2–4^. Among these, the crosstalk between SCs and cells of the immune lineages plays a pivotal role in coordinating regeneration. SCs reside in a quiescent state beneath the basal lamina of myofibers, and are defined by the expression of the paired-box transcription factor 7 (PAX7). Upon injury, SCs exit quiescence, proliferate, and differentiate into myogenic precursors under the control of myogenic regulatory factors (MRFs), including Myogenin (MYOG), to initiate and sustain the myogenic program^5^. This process is supported by infiltrating immune cells.

Neutrophils are the first responders, infiltrating the damaged tissue within 2 hours post-injury and peaking at ∼24 hours^6^. They participate in debris clearance and release pro-inflammatory cytokines and reactive oxygen species to amplify the local inflammatory response^6, 7^. This is followed by the recruitment of inflammatory monocytes, which differentiate into pro-inflammatory macrophages expressing both classical M1- and M2-associated genes (e.g., *Tnfα*, *Il-1β*, *Arg1*, *Il-10*)^8^. These cells contribute to debris clearance and promote SCs proliferation. By 3 to 4 days post-injury (dpi), a phenotypic switch occurs, leading to the emergence of anti-inflammatory macrophages that facilitate SCs differentiation and myofiber fusion^9^. The recruitment of monocytes to the injury site is dependent on CCL2/CCR2 signaling, with both resident muscle cells and macrophages contributing to chemokine expression^10–12^. In parallel, non-immune stromal cells also shape the regenerative landscape. FAPs undergo a transient expansion during the early stages of regeneration and promote SCs myogenic commitment *via* paracrine signaling^13–15^. This expansion is followed by macrophage-mediated clearance of an excess of FAPs to prevent fibrotic remodeling and re-establish homeostasis^16^. Fibroblasts also interact reciprocally with SCs to balance proliferation and fibrosis during tissue repair^17,18^.

Muscle regeneration occurs in a systemic context involving neuroendocrine regulation. Activation of the hypothalamic-pituitary-adrenal (HPA) axis in response to injury results in elevated levels of glucocorticoids (GC)^19^, that are potent anti-inflammatory steroid hormones that modulate both immune and metabolic pathways^20^. Pharmacologically, synthetic GC agonists such as dexamethasone and prednisone are widely used to treat inflammatory conditions and muscular disorders^21^. However, chronic GC exposure is associated with deleterious side effects, including muscle wasting^15, 19^. Interestingly, GC have been reported to exert context-dependent paradoxical effects on muscle physiology. While they promote terminal myogenic differentiation *in vitro* ^22^, they impair muscle regeneration *in vivo*, particularly under chronic inflammatory conditions^22–25^. These opposite effects reflect the complexity of GC signaling across different cell types. In macrophages, GCs drive the acquisition of an anti-inflammatory and pro-resolving phenotype by repressing inflammatory programs, while inducing the expression of genes associated with tissue repair. However, these effects are highly context-dependent, as GCs dynamically shape macrophage functional states according to microenvironmental signals during tissue injury and regeneration^26^.

GC effects are mediated by the glucocorticoid receptor (GR, NR3C1), a ligand-dependent nuclear receptor^27^. Upon ligand binding, GR is translocated to the nucleus where it can activate or repress gene expression. To date, more and more evidence suggests that the anti-inflammatory effects of GCs are thought to be primarily mediated by transrepression mechanisms, involving inhibition of key pro-inflammatory transcription factors such as NF-κB and AP-1, whereas many of their adverse and metabolic side effects are associated with direct transcriptional activation via glucocorticoid response elements (GREs)^28–34^. This enables context-dependent activation or repression of gene expression depending on chromatin accessibility and co-factor availability^35^.

Although the immunosuppressive functions of GC are well studied^36^, the molecular mechanisms by which GC shape macrophage homeostasis, thereby influencing muscle stem cell dynamics, remain insufficiently defined. In this study, we combine genome-wide chromatin profiling, pharmacological modulation, and *in vivo* functional analyses to uncover a myeloid-specific GC/GR regulatory circuit. We show that GC signaling via GR in macrophages controls their proliferation and polarization, which in turn governs satellite cell activation and muscle regeneration at pharmacological GC levels.

## Results

### GR in Myeloid Cells Regulates Macrophage Proliferation and Survival During Skeletal Muscle Regeneration

Analysis of GR expression from single-cell RNA-seq data obtained from regenerating tibialis anterior (TA) muscles (PRJNA577458) revealed that GR was expressed at high levels across multiple myeloid populations, including neutrophils, monocytes, and macrophages at various stages of regeneration (Fig. 1A). We therefore generated conditional knockout mice lacking GR specifically in myeloid cells (GR^Lysm-/-^ mice), in which the exon 3 of the *Nr3c1* gene (encoding GR) is selectively deleted using the *Lysm*-Cre deleter strain^37, 38^ (Fig. 1A). PCR assays on CD45⁺ / CD11B⁺ cells, isolated from TA muscles by Fluorescence-Activated Cell Sorting (FACS) 2 days after cardiotoxin (Ctx) injury (2 dpi), showed the efficient recombination of the GR floxed allele (Fig. S1B-C). Moreover, immunofluorescence analysis revealed that GR is efficiently ablated in myeloid cells of GR^Lysm-/-^ mice as exemplified for IBA1-positive cells at 2, 4 and 7 dpi (Fig. 1B).

**Figure 1:**
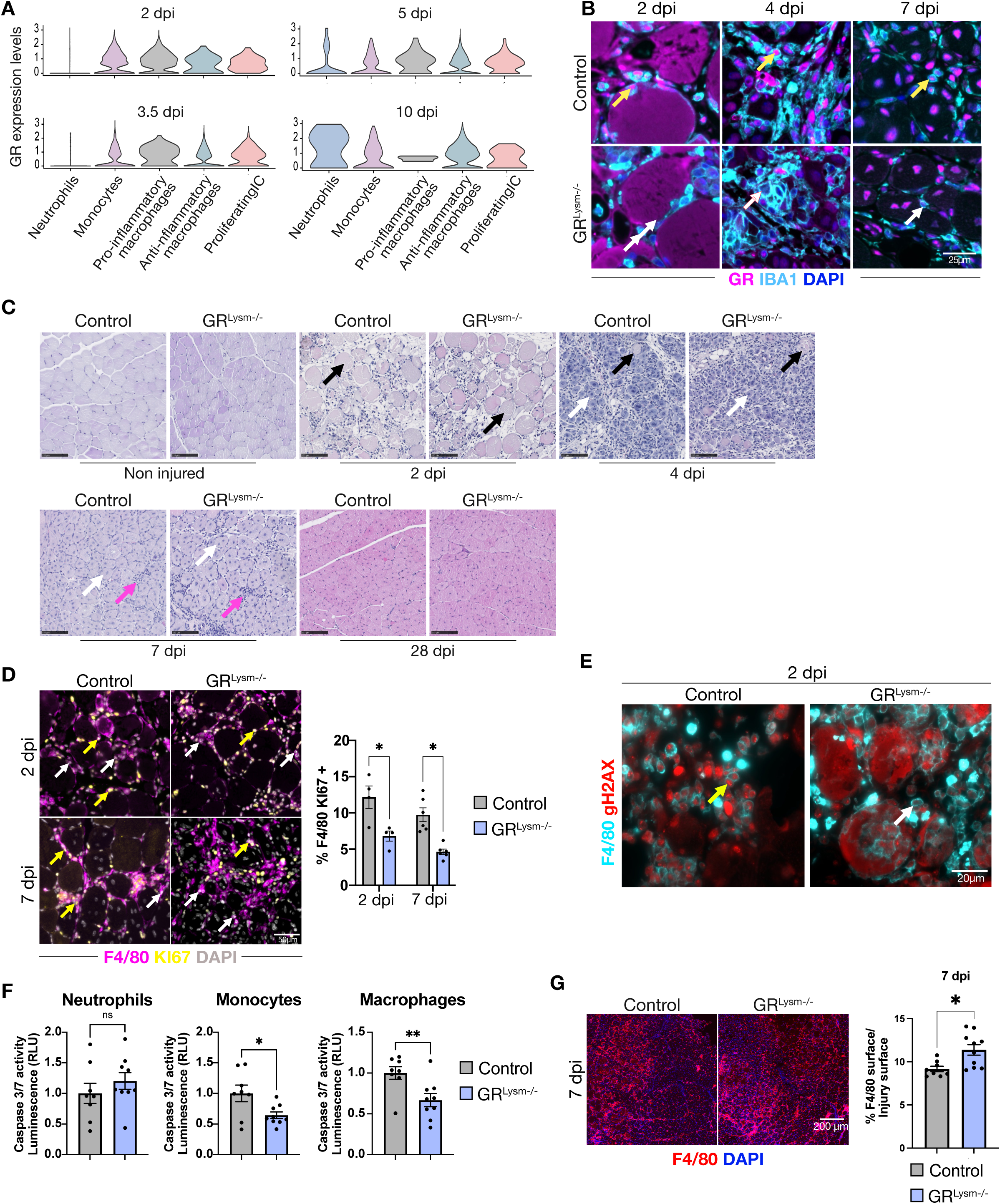
Glucocorticoid receptor deficiency in myeloid cells promotes macrophages survival and perturbs cell cycle progression. (A) Violin plots showing *Gr* expression levels from scRNA-seq analysis of injured tibialis anterior (TA) muscles of control mice at various stages of regeneration, across neutrophils, monocytes, pro-inflammatory macrophages, anti-inflammatory macrophages, and proliferating immune cells (Proliferating IC). (B) Representative immunofluorescence labeling of GR (magenta) and IBA1 (cyan) in TA muscles from control and GR^Lysm-/-^ mice at 2-, 4-, and 7-days post-injury (dpi). White arrows indicate GR-negative macrophages, yellow arrows highlight GR-positive macrophages. Nuclei were counterstained with DAPI. Scale bar, 25 μm. (C) Representative hematoxylin and eosin staining of TA muscles of control and GR^Lysm-/-^ mice under non-injured condition, and at 2, 4 and 7 dpi. Scale bar, 100 µm. (D) Representative immunofluorescent detection of F4/80 (magenta), and KI67 (yellow) and corresponding quantifications in TA of control and GR^Lysm-/-^ mice 2 and 7 dpi. Nuclei were stained with DAPI. Scale bar, 50 μm. Data are presented as mean ± SEM. Statistical test used was Student’s t-test; * = p < 0.05. (E) Representative immunofluorescent detection of F4/80 (cyan), and gH2AX (red) in TA of control and GR^Lysm-/-^ mice 2 and 7 dpi. Yellow arrows indicate gH2AX-positive macrophages while white arrows indicate gH2AX-negative macrophages. Scale bar, 20 μm. (F) Caspase 3/7 activity measured in FACS-purified control and GR^Lysm-/-^ neutrophils, monocytes and macrophages 2 dpi. RLU, relative luminescence units. Data are presented as mean ± SEM. Statistical test used was Student’s t-test; ns, non-significant; * = p < 0.05. (G) Representative immunofluorescence staining of F4/80 (red) and quantification of the area occupied by F4/80-positive cells relative to the injury area in TA muscles from control and GR^Lysm-/-^ mice at 2, 4, and 7 dpi. Statistical test used was Student’s t-test; ns, non-significant; * = p < 0.05. Scale bar, 200 µm.

At 2 dpi, a time point at which all major myeloid populations are present, muscle histology assessed by Hematoxylin and Eosin staining (H&E) was similar between control and GR^Lysm-/-^, with abundant number of necrotic fibers and immune infiltration, compared to a non-injured muscle (Fig. 1C). As GR is known to be involved in cell cycle progression, we examined the proliferation and apoptosis of control and GR deficient myeloid cells. Importantly, KI67 and F4/80 co-immunostaining showed that the percentage of proliferating macrophages was decreased by more than two-fold upon GR loss from 2 dpi. (Fig. 1D). We next evaluated whether GR impacts cell survival by performing co-staining with antibodies directed against γH2AX, that stands for DNA damages, and the macrophage markers F4/80, and by measuring caspase 3/7 activity in FACS-isolated myeloid cells. We noticed that fewer γH2AX-positive macrophages were detectable in TA of GR^Lysm-/-^ mice 2 dpi compared to controls (Fig. 1E), showing that GR is required for double-strand DNA break formation. Moreover, at this regeneration stage, caspase activity was significantly lower in macrophages and monocytes lacking GR, but remained similar between control and mutant neutrophils (Fig. 1F).

We next investigated the consequences of altered monocytes and macrophages cell cycle progression. While histological kinetics analysis by H&E staining assessed 4 to 28 dpi did not reveal major architectural differences between regenerating myofibers of control and mutant mice, a marked increase in immune cell infiltration was observed in GR^Lysm-/-^ muscles at 7 dpi (Fig. 1C, 1D and S1D). Immunostaining for the macrophage marker F4/80 revealed that this infiltration was at least in part due to macrophages despite decreased proliferation rate (Fig. 1D and 1G). Together our data show that GR is required for monocytes and macrophages cell cycle progression and survival associated with macrophage accumulation at 7 dpi.

### Myeloid GR Deficiency Has Moderate Impact on Satellite Cell and Fibro-Adipogenic Progenitor Dynamics

Since macrophages are known to influence the proliferation and differentiation of both muscle stem cells and FAPs upon injury^39^, we investigated the impact of increased macrophage accumulation observed in GR^Lysm-/-^ mice on myogenesis and fibrosis. 7 dpi, the number of PAX7-positive satellite cells (SCs) was comparable between control and GR^Lysm-/-^ mice, as was SCs proliferation, revealed by PAX7 and KI67 co-staining (Fig. 2A). Furthermore, muscle stem cell differentiation, assessed by MYOG expression, was similar between genotypes, including the percentage of MYOG-positive proliferating cells (Fig. 2A). Consistently, the number of centro-nucleated fibers was not affected by GR loss in myeloid cells at 7 dpi (Fig. 2B). These data show that GR invalidation in myeloid cells does not impact the proliferation or differentiation of SCs at endogenous GC levels. In addition, immunofluorescent detection of alpha-actinin (ACTN1) did not reveal gross abnormalities in muscle fiber structure (Fig. 2C). Terminal differentiation, assessed by the quantification of TA myofiber cross-sectional area at 28 dpi, was similar between GR^Lysm-/-^ mice and their control littermates (Fig. 2D). Consistently, no differences in muscle weight were observed between genotypes at 4, 7, 15 and 21 dpi (Fig. S1E), showing that GR signaling in myeloid cells does not impact muscle growth during muscle regeneration in physiological conditions.

**Figure 2:**
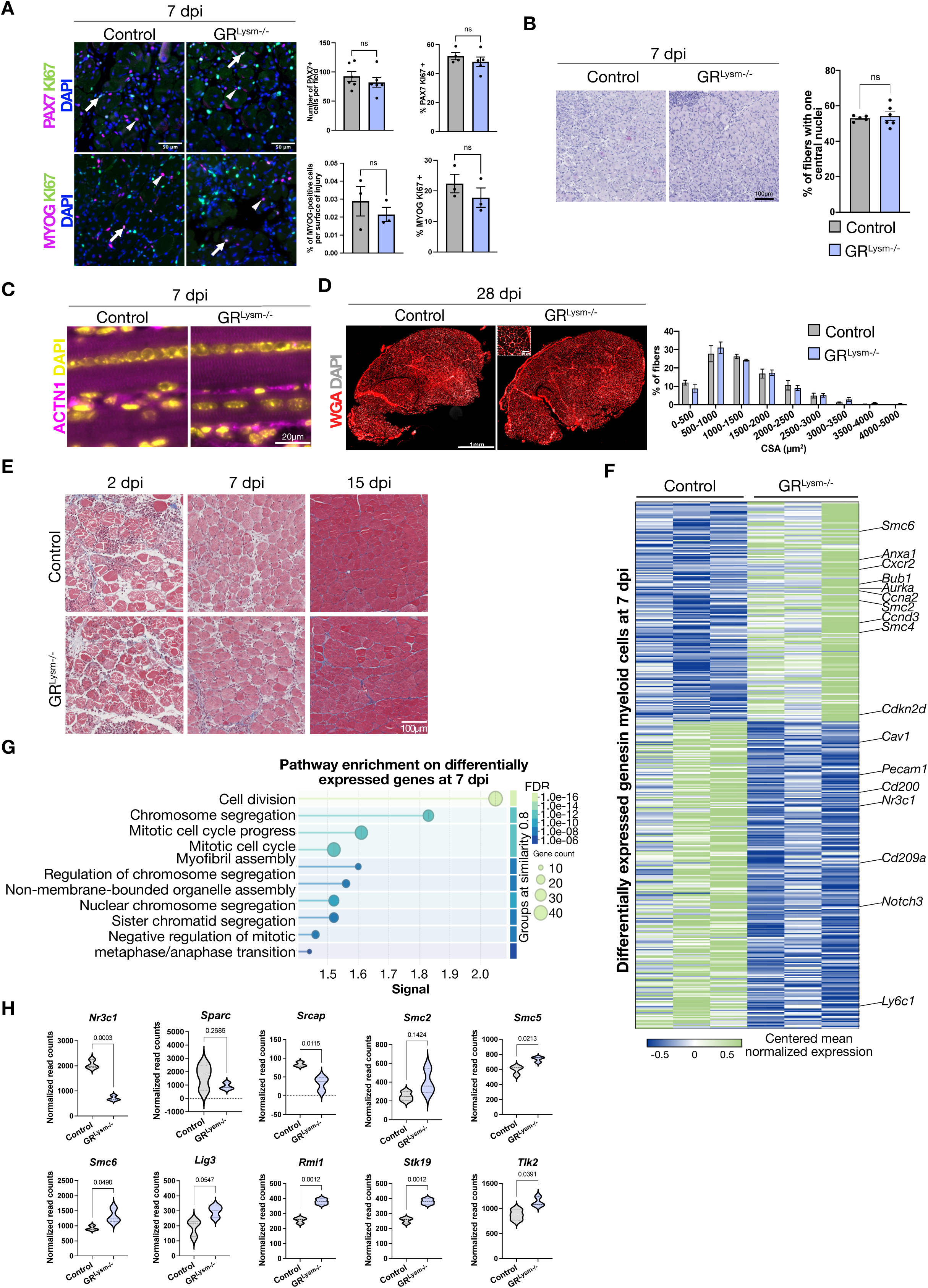
Glucocorticoid receptor deficiency in myeloid cells does not impact muscle regeneration. (A) Representative immunofluorescent detection of PAX7 or MYOG (magenta), and KI67 (green) and corresponding quantifications in TA of control and GR^Lysm-/-^ mice 7 dpi. Nuclei were stained with DAPI. Scale bars, 50 μm. Data are presented as mean ± SEM. Statistical test used was Student’s t-test; ns, non-significant. (B) Representative hematoxylin and eosin staining of 7 days-injured TA muscles from control and GR^Lysm-/-^ mice. Quantifications shows the percentage of regenerative fibers containing a single nucleus. Scale bar, 250 µm. (C) Representative immunofluorescent detection of ACTN1 (magenta) in TA of control and GR^Lysm-/-^ mice 7 dpi. Nuclei were stained with DAPI (yellow). Scale bar, 20 µm. (D) Representative staining of Wheat Germ Agglutinin (WGA, red) in 28-days-injured TA muscles of control and GR^Lysm-/-^ mice, and corresponding fiber cross-sectional area (CSA) distribution across the injury site. No significant difference was observed in fiber size distribution between genotypes. Nuclei were stained with DAPI (gray). Scale bars, 1 mm (main image) and 50 μm (magnified inset). (E) Representative Masson trichrome staining of TA of control and GR^Lysm-/-^ mice at indicated time points of muscle regeneration. Scale bar, 15 µm. (F) Heatmap showing the mean-centered normalized expression of selected genes from bulk RNA-sequencing of FACS-sorted CD11b⁺ cells isolated from TA muscles of control and GR^Lysm-/-^ mice 7 dpi. (G) Pathway analysis of down- and up-regulated genes in 7-days-injured TA of control and GR^Lysm-/-^ mice. (H) Violin plots showing normalized read counts of selected genes that were differentially expressed in bulk RNA-sequencing of FACS-sorted CD11b⁺ cells isolated from TA muscles of control and GR^Lysm-/-^ mice 7 dpi.

Moreover, the number of adipocytes was similar between GR^Lysm-/-^ and control mice across the regeneration timeline, as exemplified at 21 dpi (Fig. S1F), and Gomori Trichrome staining combined with immunofluorescent detection of collagen IV (COL4) showed comparable levels of fibrosis between groups 2, 7, and 15 dpi (Fig.2E and Fig. S1G). Altogether, our findings show that despite GR loss-induced perturbation in macrophage dynamics, muscle stem cell differentiation and FAPs fate decisions remain largely unaffected, indicating that GR deficient myeloid cells have only a moderate influence on the regenerative microenvironment at endogenous GC levels.

### Transcriptomic Profiling Reveals GR-Dependent Gene Networks in Myeloid Cells

To delineate the GR-dependent transcriptional signature in myeloid cells, we performed transcriptomic profiling on FACS-isolated CD11B⁺ cells from injured TA at 7 dpi (Fig. 2F). Among the differentially expressed genes, 299 were up- and 369 were downregulated in the absence of GR. Over-representation analysis using the Gene Ontology Biological Process database revealed that differentially expressed genes were enriched in pathways related to regulation of cell proliferation and genome organization (Fig. 2G). Among those genes, the expression of *Sparc* that is involved in G1/G0 phase regulation and apoptosis^40^ and *Srcap*, that contributes to chromatin remodeling was reduced by two-fold, while that of *Smc2, Smc5, Smc6* ATPases from the cohesin-dependent complexes was increased by ∼1.5-fold, as well as additional chromatin-associated factors (Fig. 2H and S1H). Notably, several genes involved in DNA repair and genome stability, including *Lig3* and *Rmi1*, as well as the nuclear kinase *Stk19* and the cell-cycle regulator *Tlk2*, displayed marked elevated expression in GR^Lysm-/-^ muscles related control littermates (Fig. S1H). Together, these transcriptional alterations highlight that GR positively or negatively regulates the expression of genes controlling chromosome dynamics and cell-cycle progression.

To identify genes potentially directly regulated by GR underlying the altered cell cycle dynamics observed upon GR loss, we next characterized GR cistrome by CUT&RUN on macrophages isolated from injured TA muscles. Our experiment uncovered 7,464 GR binding sites, primarily located in intronic and intergenic regions, with an average distance of ±50 kb from the TSS (Fig. S2A-B), indicating that GR is recruited to distal enhancers, in agreement with previous reports (GSE93736)^41^ (Fig. S2C-D). To determine the chromatin landscape relative to GR binding sites, we assessed the genomic profile of H3K4me3 and H3K27ac, that are hallmarks for active promoters and enhancers, respectively. CUT&RUN analyses unveiled 17,202 H3K4me3 peaks predominantly located within ±2 kb of transcription start sites (TSS), and 15,768 H3K27ac peaks distributed across promoter and enhancer regions within ±50 kb of TSS (Fig. S2A-B). Notably, nearly 70% of genes with read counts >100 in our transcriptome dataset showed H3K4me3 enrichment, supporting the robustness of our profiling (Fig. S2E), and GR binding overlapped with active chromatin regions marked by H3K4me3 and H3K27ac, indicating that GR is predominantly recruited at transcriptionally permissive loci (Fig. 3A-B). *De novo* motif analysis revealed that GR is bound to 5ʹ-AGAVCAgccTGGnnn-3ʹ glucocorticoid response elements (GREs) in ∼45% of targeted regions (Fig. 3C). Pathway enrichment analysis on the 5,133 GR-bound genes highlighted biological processes related to efferocytosis and RAP1 signaling, as exemplified Fig 3B, D but did not identify cell cycle-related pathways, even though transcriptome and histological analyses converged toward alterations of macrophage cell cycle dynamics (Fig. 1D-F and Fig. 2F-H). Instead, Gene Ontology analysis of genes associated with GR binding sites highlighted terms associated with developmental processes and cellular responses, including “nervous system development”, “cellular response to organic substance”, “regulation of signaling”, and “neurogenesis”. To further refine our analysis, we intersected GR-bound genes with differentially expressed genes identified by RNA-seq, yielding only 124 common genes, related to extracellular matrix organization, regulation of cell growth, and developmental processes (Fig. 3E-F), indicating that cell cycle defects observed upon GR deletion are unlikely to be driven by direct transcriptional regulation.

**Figure 3:**
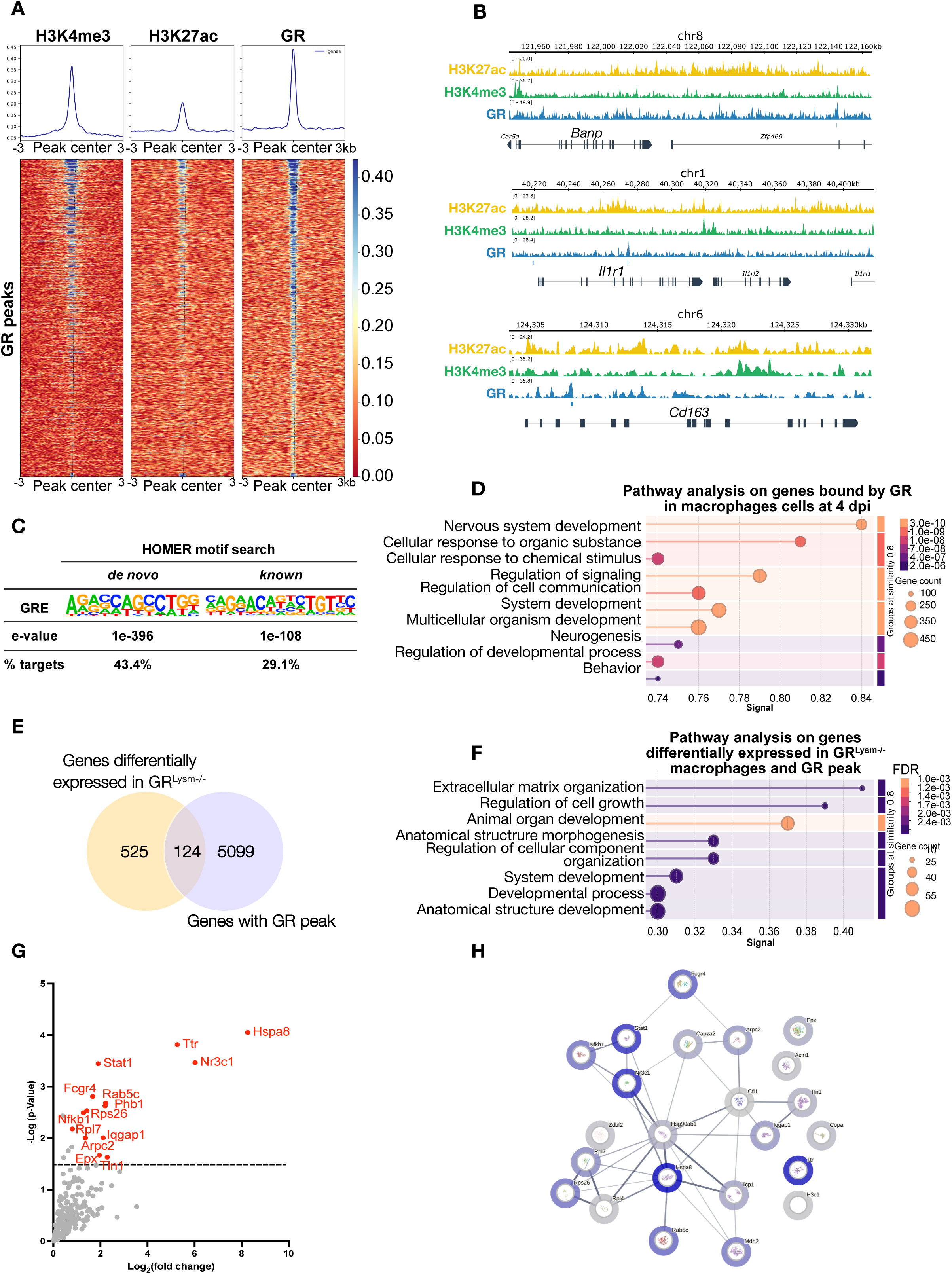
Glucocorticoid receptor cistrome and proteomic analyses in macrophages. (A) Tag density map of H3K4me3, H3K27ac and GR in myeloid cells, +/- 5 kb from the GR peak center sorted by GR peak length, and corresponding average tag density profiles. (B) Localization of GR, H3K4me3 and H3K27ac at representative promoter and intergenic regions on the chromatin of myeloid cells by CUT&RUN. Enrichment score showing the levels of confidence of published datasets is depicted as a heatmap. (C) HOMER motif analysis of GR binding sites located at intergenic, intronic or promoter regions. p-value: hypergeometric testing. (D) Pathway analysis performed on GR common target genes identified in macrophages of TA of control and GR^Lysm-/-^ mice 4 dpi. (E) Overlap of the genes bound by GR in macrophages and genes differentially expressed in bulk RNA-sequencing of FACS-sorted CD11b⁺ cells isolated from TA muscles of control and GR^Lysm-/-^ mice 7 dpi. (F) Pathway analysis performed on GR common target genes identified in macrophages of TA of control and GR^Lysm-/-^ mice 4 dpi and differentially expressed in bulk RNA-sequencing of FACS-sorted CD11b⁺ cells isolated from TA muscles of control and GR^Lysm-/-^ mice 7 dpi. (G) Volcano plot depicting GR partners identified by mass-spectrometry in FACS-isolated myeloid cell extracts 4 dpi. (H) STRING analysis of the nuclear proteins co-immunoprecipitated with GR in FACS-isolated myeloid cell ex tracts. Line thickness indicates the strength of data support for protein-protein interactions 4 dpi.

To explore this latter possibility, we performed GR immunoprecipitation followed by mass spectrometry (LC-MS/MS) to identify proteins interacting with GR, aiming to uncover potential mechanisms by which GR may regulate macrophage cell cycle dynamics independently from direct DNA binding. GR interactome uncovered 23 interacting proteins, from which GR (NR3C1) was the top identified protein, thereby validating the specificity for our approach. Among these newly identified GR partners, only 6 were previously identified according to BioGrid, namely HSP90AB1, HSPA8, NFKB1, NR3C1, RPS26 and ZDBF2 (Table S3). Pathway analysis on GR-interacting proteins revealed that they mainly belong to immunity (e.g. STAT1 and NFKB), but also uncovered chaperones (Fig. 3G). Notably, the Heat Shock Protein Family A (Hsp70) Member 8, HSPA8 (also called heat shock cognate 71 kDa protein or Hsc70) was the most enriched peptide, which would be in favor of an inactive, low-affinity GR conformation. In contrast, association to HSP90beta (referred as HSP90AB1) was lower, suggesting limited formation of the mature high-affinity complexes in macrophages^42^.

Beyond these proteins, GR partners were involved in diverse cellular processes such as cytoskeletal organization (TLN1, IQGAP1, ARPC2), intracellular trafficking (RAB5C), RNA and ribosome biology (RPS26, RPL7), extracellular matrix regulation (COL2A1), and immune effector functions (EPX, FCGR4, NRDC, TTR, PRSS3B, TRY10). We also identified PHB1 (Prohibitin-1), a co-regulator previously implicated in the modulation of nuclear receptor signaling^43^, suggesting a potential role in fine-tuning GR activity in myeloid cells.

Altogether, our findings show that GR does not directly regulate canonical cell cycle genes in macrophages, but rather exerts its effects through indirect transcriptional programs and protein interaction networks, supporting a model in which GR integrates chromatin-dependent and non-genomic mechanisms to regulate macrophage function during muscle regeneration.

### Dexamethasone Administration During the Anti-Inflammatory Phase Has Limited Impact on Muscle Regeneration

To test whether GR activity is affected by exposure to synthetic agonists, we next aimed at investigating GR molecular functions at pharmacological GC levels. To explore the functional role of GC signaling during the resolution phase of muscle regeneration, we intraperitoneally administered dexamethasone (Dex) to control and GR^Lysm-/-^ at 4, 5 and 6 dpi (Fig. 4A). Histological analysis performed at 7 dpi did not reveal major influence on muscle regeneration for both genotypes (Fig. 4B). The extent of myofiber necrosis was similar between the various conditions, and immune cell infiltration was not markedly altered (Fig. 4B). Interestingly, Dex induced the formation of vesicle-like structures at the periphery of regenerating myofibers in a myeloid GR-independent manner (Fig. 4B). These vesicles were membrane-less and did not stain for the lipid droplet marker perilipin, suggesting they are not classical lipid accumulations.

**Figure 4:**
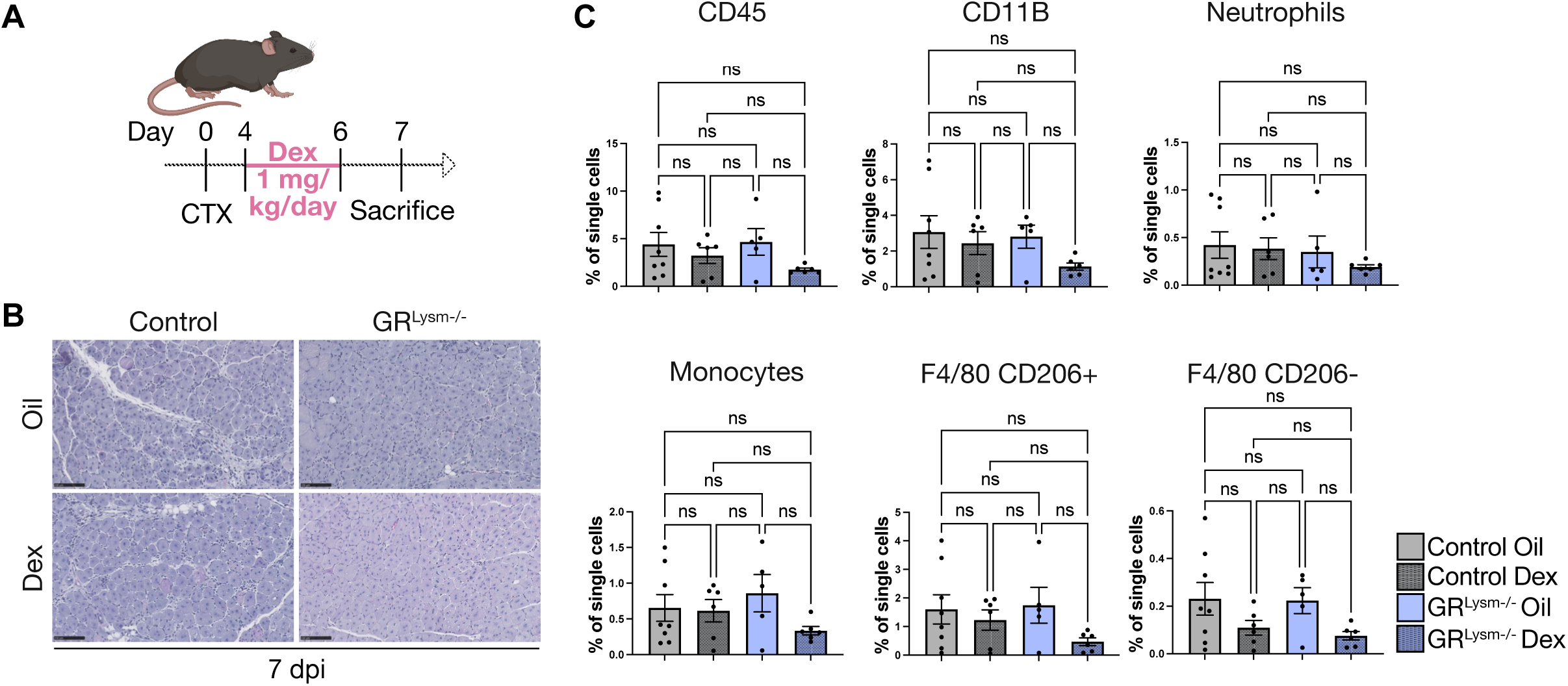
Dexamethasone treatment during the anti-inflammatory phase does not affect muscle regeneration or myeloid cell composition. (A) Schematic representation of the treatment regimen for control and GR^Lysm-/-^ mice. (B) Representative hematoxylin and eosin staining of tibialis muscles (TA) of dexamethasone-(Dex) or vehicle-treated (Oil) control and GR^Lysm-/-^ mice 7 days after injury. Scale bar, 100 µm. (C) Percentage of myeloid cells in 7-days-injured TA of Dex- or Oil-treated control and GR^Lysm-/^. Data are presented as mean ± SEM. Statistical test used is Two-way ANOVA with Tukey *post-hoc* correction; ns, non-significant; ** = p < 0.01; *** = p < 0.001.

Moreover, flow cytometry analysis revealed no significant differences in the proportions of CD45⁺ and CD11b⁺ cells, as well as in neutrophil, monocyte, and macrophage (CD206^+^ and CD206^-^) populations across the four analyzed conditions (Fig. 4C). Altogether, these findings show that short-term activation of GR signaling during the anti-inflammatory phase has only minor effects on the immune landscape and does not significantly improve or impair muscle recovery.

### Dexamethasone Administration During the Pro-Inflammatory Phase Impairs Muscle Regeneration in a Myeloid GR-Dependent Manner

To investigate the consequences of an early GC treatment on muscle regeneration, Dex was intraperitoneally injected during the acute inflammatory phase at 0, 1 and 2 dpi (Fig. 5A). Histological examination at 2 dpi revealed no major differences between vehicle- and Dex-treated control and GR^Lysm-/-^ mice, the four groups displaying extensive myofiber necrosis and stromal reaction (Fig. S3A). However, Dex-treated control and GR^Lysm-/-^ mice exhibited a high number of persistent necrotic fibers, reminiscent of the early damage phase, related to vehicle-treated cohorts by 4 dpi, despite the emergence of nascent myofibers (Fig. S3A-B). By 7 dpi, immunofluorescence and histological analyses revealed that even though inflammation was partially resolved in vehicle-treated mice, TA muscles of Dex-treated control and GR^Lysm-/-^ individuals displayed sustained necrosis and an exacerbated stromal response (Fig. 5B-C), indicating that Dex delays muscle repair.

**Figure 5:**
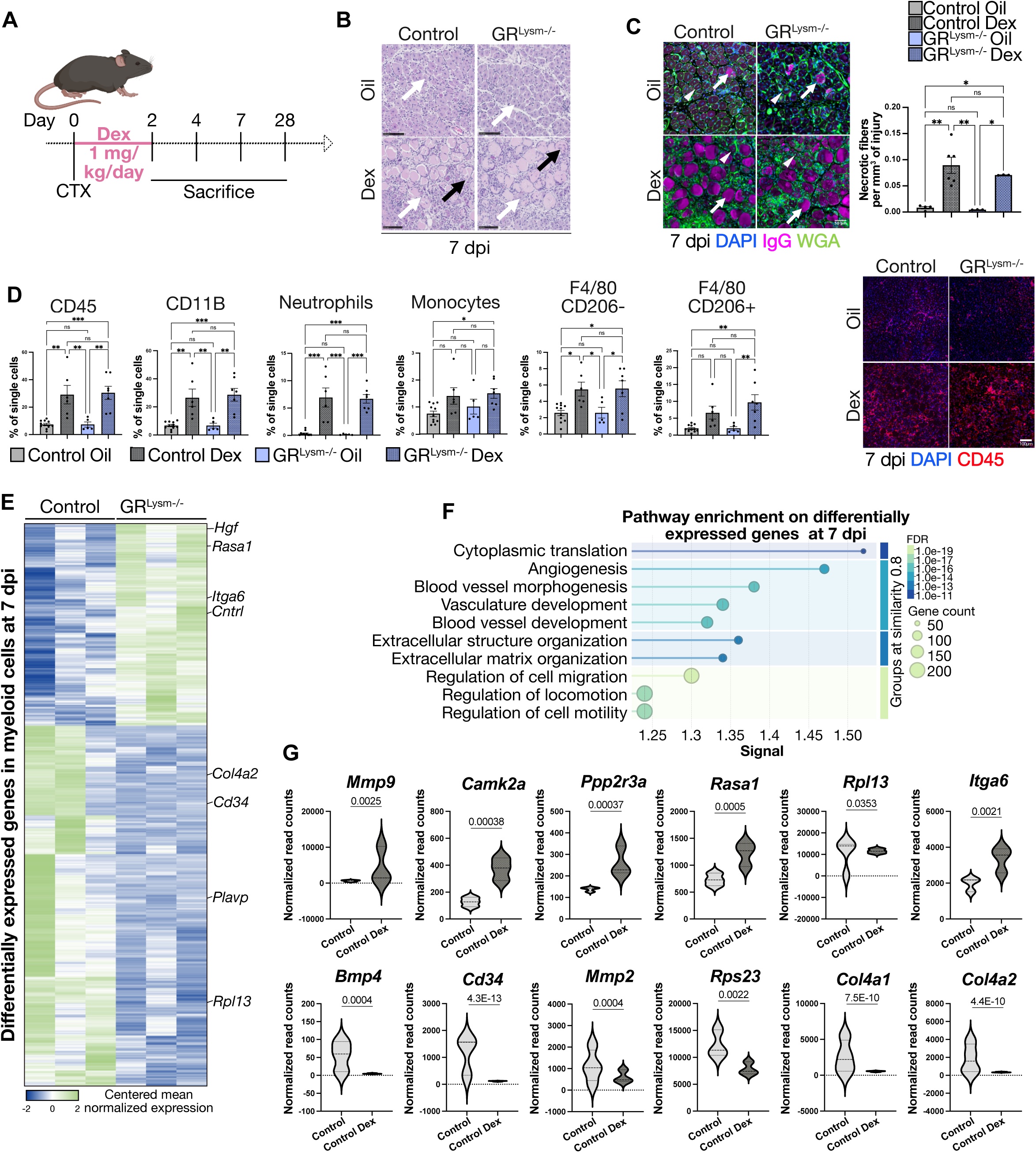
Dexamethasone treatment during the pro-inflammatory phase impairs muscle regeneration. (A) Schematic representation of the treatment regimen for control and GR^Lysm-/-^ mice. (B) Representative hematoxylin and eosin staining of tibialis muscles (TA) of Dex- or vehicle-treated (Oil) control and GR^Lysm-/-^ mice 7 days after injury. Scale bar, 100 µm. (C) Representative immunofluorescence staining of Wheat Germ Agglutinin (WGA, green) and IgG (magenta) in TA muscles of Dex- or Oil-treated control and GR^Lysm-/-^ mice, 7 days after injury. Quantification represents the percentage of necrotic (IgG⁺) fibers relative to the injury surface. Nuclei were stained with DAPI. Scale bar, 50 µm. Data are presented as mean ± SEM. Statistical test used is Two-way ANOVA with Tukey *post-hoc* correction; ns, non-significant; ** = p < 0.01; *** = p < 0.001. (D) Percentage of myeloid cells and representative immunofluorescent detection of CD45 (red) in 7-days-injured TA of Dex- or Oil-treated control and GR^Lysm-/-^. Nuclei were stained with DAPI. Data are presented as mean ± SEM. Statistical test used is Two-way ANOVA with Tukey *post-hoc* correction; ns, non-significant; ** = p < 0.01; *** = p < 0.001. (E) Heatmap showing mean-centered normalized expression of selected genes from bulk RNA-sequencing of FACS-sorted CD11b⁺ cells isolated from 7-days-injured TA muscles of control Dex- or Oil-treated. (F) Pathway analysis of down- and up-regulated genes in 7-days-injured TA of control Dex- or Oil-treated. (G) Violin plots showing normalized read counts of selected genes that were significantly differentially expressed in bulk RNA-sequencing of FACS-sorted CD11b⁺ cells isolated from 7-days-injured TA muscles of control Dex- or Oil-treated.

Given the potent GC anti-inflammatory properties, we next assessed their impact on immune cell dynamics *via* flow cytometry on injured TA, 2, 4, and 7 dpi (Fig. S3C). Our data revealed that the number of CD45⁺ leukocytes and CD11b⁺ myeloid cells was similar upon Dex-treatment in both control and GR^Lysm-/-^ mice 2 dpi (Fig. S3D), as well as that of neutrophils and monocytes, that are predominant at this stage comprising ∼8% and ∼15% of total live cells, respectively. In contrast, CD206^-^ and CD206^+^ cells within the F4/80⁺ macrophage population were decreased, suggesting that an early GC treatment delays monocyte-to-macrophage transition (Fig. S3D).

At 4 dpi, total leukocyte and myeloid cell frequencies (∼30% of living cells) were comparable between vehicle- and Dex-treated control and GR^Lysm-/-^ animals (Fig. S3E). While monocyte and macrophage number was not affected by Dex treatment at this time point, neutrophil fraction was augmented by ∼3-folds in Dex-treated control mice, relative to vehicle-treated individuals, most probably due to the presence of persistent necrotic fibers. This induction was reduced in GR^Lysm-/-^ animals, indicating that additional neutrophil recruitment 4 dpi is myeloid GR-dependent.

Notably, while immune cell clearance was evident in vehicle-treated mice 7 dpi, with leukocytes representing <10% of total living cells, Dex-treated littermates showed persistent inflammation, with ∼35% of CD45⁺ cells (Fig. 5D), in agreement with our histological analyses. Myeloid cell numbers remained elevated across all subpopulations, including a marked accumulation of CD206⁺ macrophages. Neutrophil percentage was even more increased than at 4 dpi, reaching levels comparable to those observed at 2 dpi, indicating that delayed regeneration may involve the re-initiation of an inflammatory response. This second wave of myeloid infiltration was not affected by GR deletion in myeloid cells, suggesting GR-independent mechanisms of recruitment.

To gain further insight into the molecular pathways regulated by pharmacological glucocorticoids in myeloid cells, we performed RNA-seq analysis on myeloid cells isolated by FACS from 7 dpi regenerating muscles of control mice treated by Dex or vehicle. Transcriptomic analysis identified 754 upregulated and 1337 downregulated genes following Dex treatment of control individuals (Fig. 5E). Gene Ontology pathway analysis revealed an enrichment in biological processes related to cell motility and vascular remodeling (Fig. 5F), including *Mmp9, Camk2a, Kif20b* and *Ppp2r3a* that contribute to cell motility, *Has1*, *Wnt4*, *Mmp3* and *Selp* involved in cell adhesion, *Rasa1*, *Nrp2*, *Tmem100*, *Bmp4* and *Cd34* related to angiogenesis, *Col1a1*, *Col3a1*, *Fbn1*, *Fn1*, *Mmp2* and *Postn* associated with extracellular matrix, as well as *Dusp6, Smad7, Spry2, Rasa1* and *Itga6* implicated in the negative regulation of cell migration. Together, our data show that pharmacological GC strongly impact muscle regeneration by primarily altering transcriptional programs governing cell dynamics.

### Synthetic Glucocorticoid Treatment Reveals an Intrinsic GR-mediated Crosstalk between Macrophages and Satellite Cells

Given the elevated macrophage numbers at 7 dpi, we next assessed their proliferative capacity. Dex treatment from 0 to 2 dpi doubled macrophage proliferation in control but not in GR^Lysm-/-^ mice (Fig. 6A). Notably, the CD163^+^ macrophage subset was KI67-negative in either control or Dex-treated conditions (Fig. 6B), thus showing that three-day Dex treatment induces recruited macrophage proliferation in a GR-dependent manner.

**Figure 6:**
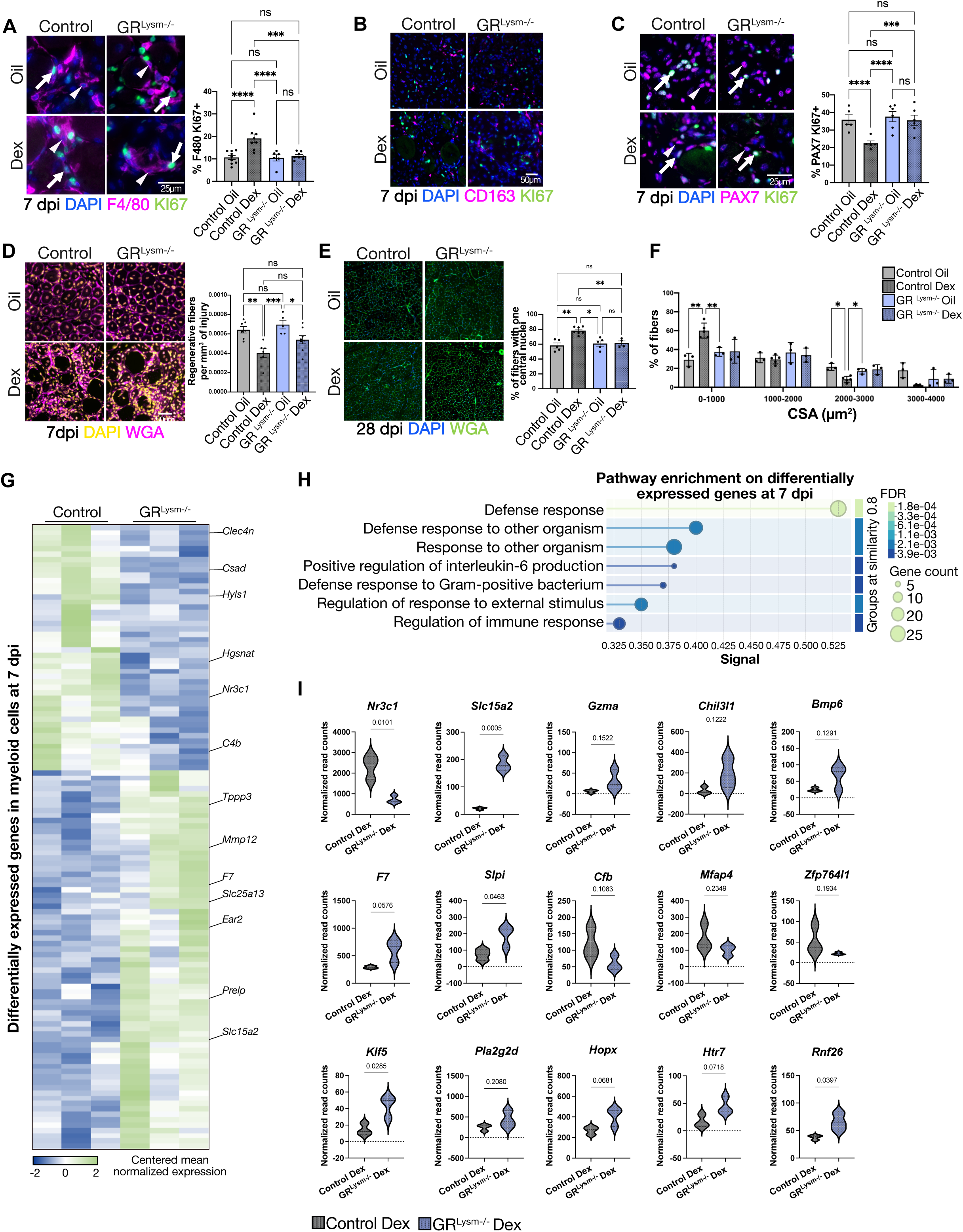
Dexamethasone impairs myogenesis during the pro-inflammatory phase through GR-dependent signaling in myeloid cells. (A) Representative immunofluorescent detection of F4/80 (magenta), and KI67 (green) and corresponding quantifications in 7-days injured TA of Dex- or Oil-treated control and GR^Lysm-/-^mice. Nuclei were stained with DAPI. Scale bars, 25 μm. Data are presented as mean ± SEM. Statistical test used is Two-way ANOVA with Tukey *post-hoc* correction; ns, non-significant; ***=p <0.001; **** = p < 0.0001. (B) Representative immunofluorescent detection of CD163 (magenta) and KI67 (green) in 7-days injured TA of Dex- or Oil-treated control and GR^Lysm-/-^mice, only few, if any, CD163⁺ cells were positive for KI67 in any condition. Nuclei were stained with DAPI. Scale bars, 50 μm. (C) Representative immunofluorescent detection of PAX7 (magenta), and KI67 (green) and corresponding quantifications in 7-days injured TA of Dex- or Oil-treated control and GR^Lysm-/-^mice. Nuclei were stained with DAPI. Scale bars, 25 μm. Data are presented as mean ± SEM. Statistical test used is Two-way ANOVA with Tukey *post-hoc* correction; ns, non-significant; ***=p <0.001; **** = p < 0.0001. (D) Representative immunofluorescence staining of Wheat Germ Agglutinin (WGA, maganta) in TA muscles of Dex- or Oil-treated control and GR^Lysm-/-^ mice, 7 days after injury. Quantification represents the percentage of regenerative fibers relative to the injury surface. Nuclei were stained with DAPI (yellow). Scale bar, 50 µm. Data are presented as mean ± SEM. Statistical test used is Two-way ANOVA with Tukey *post-hoc* correction; ns, non-significant; * = p < 0.05; ** = p < 0.01; ***=p <0.001. (E) Representative immunofluorescence staining of Wheat Germ Agglutinin (WGA, green) in TA muscles of Dex- or Oil-treated control and GR^Lysm-/-^ mice, 28 days after injury. Quantification shows the percentage of regenerative fibers containing a single nucleus (F) and the corresponding fiber cross-sectional area (CSA) distribution within the injury site. Nuclei were stained with DAPI. Data are presented as mean ± SEM. Statistical test used is Two-way ANOVA with Tukey *post-hoc* correction; ns, non-significant; * = p < 0.05; ** = p < 0.01. (G) Heatmap showing mean-centered normalized expression of selected genes from bulk RNA-sequencing of FACS-sorted CD11b⁺ cells isolated from 7-days-injured TA muscles of control and GR^Lysm-/-^ mice Dex-treated. (H) Pathway analysis of down- and up-regulated genes in 7-days-injured TA muscles of control and GR^Lysm-/-^ mice Dex-treated. (I) Violin plots showing normalized read counts of selected genes that were significantly differentially expressed in bulk RNA-sequencing of FACS-sorted CD11b⁺ cells isolated from 7-days-injured TA muscles of control and GR^Lysm-/-^ mice Dex-treated.

To determine if Dex-associated increase in myeloid cell number and proliferation affects muscle stem cell homeostasis, we performed a KI67⁺/PAX7⁺ co-immunostaining. Our data revealed that Dex treatment significantly reduced satellite cell proliferation by ∼1/3, and that this effect was dependent on GR in myeloid cells (Fig. 6C). These findings indicate that GR-dependent signals from myeloid cells suppress satellite cell proliferation in response to early Dex treatment. Moreover, immunofluorescence revealed a substantial reduction in regenerative myofibers number at 7 dpi (1.5-fold) in Dex-treated mice, but not at 4 dpi (Fig. 6D, S3F). Long-term analysis at 28 dpi showed an increase in the number of fibers with only one central nucleus, indicative of ongoing regeneration, in both control Dex-treated mice, and this phenotype was abolished in GR^Lysm-/-^ mice (Fig. 6E, S3A). Likewise, the Dex-induced reduction in myofiber cross-sectional area was not GR-dependent (Fig. 6F), showing that GR in myeloid cells affects myogenesis and myofiber hypertrophy.

To determine the molecular mechanisms through which Dex treatment exclusively in control mice influences macrophage proliferation, satellite cell activity, and muscle hypertrophy, we analyzed the genes differentially expressed between control and mutant mice treated with Dex. This analysis identified 72 upregulated and 47 downregulated genes (Fig. 6G). Gene Ontology enrichment revealed a marked overrepresentation of immune-related categories, including “defense response,” “response to other organism,” “regulation of immune response,” and “defense response to Gram-positive bacterium.” These processes encompassed robustly induced genes such as *Slc15a2, Gzma, Cd36, Bmp6, F7, Chi3l1*, and *Slpi*, which are associated with antimicrobial activity, lipid metabolism, and inflammatory activation. Conversely, several genes involved in regulation and structural organization, including *CV, Mfap4, Myo1b,* and *Zfp764l1*, were consistently repressed. Additional enrichment for “regulation of response to external stimulus” and “positive regulation of interleukin-6 production” was observed, driven by genes such *as Klf5, Pla2g2d, Hopx, Htr7*, and *Rnf26*, indicating a selective modulation of cytokine signaling and stress adaptation (Fig. 6H-I).

Together, these results demonstrate that early GC exposure disrupts the macrophage inflammatory-to-anti-inflammatory transition in both GR-dependent and independent manners. GR activation in myeloid cells promotes macrophage proliferation and impairs their phenotype, henceforth skewing satellite cell proliferation and muscle regeneration. This establishes a crosstalk between myeloid cells and satellite cells, tightly regulated by myeloid GR during early repair.

## Discussion

Understanding the fundamental role of glucocorticoids during muscle repair is a key therapeutic question. While the glucocorticoid receptor is robustly expressed across myeloid populations recruited following injury, its deletion in these cells does not markedly impair the overall outcome of muscle repair at physiological glucocorticoid levels. Indeed, satellite cell proliferation and differentiation, fibro-adipogenic progenitor fate, and long-term muscle growth remain largely unaffected. These findings indicate that myeloid GR is dispensable for the execution of the core myogenic program under homeostatic regenerative conditions.

In contrast, GR deficiency profoundly alters macrophage dynamics. GR is well-known to control mitotic progression and chromosome segregation^44^. Here we observe an increased number of macrophages despite a marked reduction in their proliferative capacity, indicating that macrophage accumulation is uncoupled from local proliferation. Instead, our data point toward enhanced survival, supported by reduced apoptotic activity and decreased DNA damage in GR-deficient monocytes and macrophages. These findings are consistent with previous studies showing that GR effects on myeloid cell proliferation and survival can vary according to the experimental settings. Indeed, while glucocorticoids effects are mainly anti-proliferative in lymphoid cells, their actions in myeloid populations are more heterogeneous, and can either inhibit and promote proliferation depending on the activation state and microenvironment (^45–48^ and references within). In macrophages, results indicate that GR acts as a sensor of inflammatory and environmental signals rather than a direct suppressor of proliferation, thus contributing to the balance between cell cycle progression and survival. Similarly, glucocorticoids have been shown to regulate survival pathways differently depending on the type of immune cell, thus confirming the cell-specific role of GR in controlling the fate of myeloid cells (^49^ and references within). Consistent with a role of GR in the maintenance of genome integrity, we observed a marked reduction in γH2AX staining in GR deficient myeloid cells. γH2AX is a well-established marker of DNA double strand breaks involved in DNA damage response. Previous studies have shown that physiological concentrations of glucocorticoids induce DNA double strand breaks in a GR dependent manner, and that GR orchestrates cell cycle regulation in collaboration with the ARID1a/P53BP1 complex^50^. In this context, the reduced γH2AX signal observed in the absence of GR could reflect either a decrease in DNA damage accumulation or perturbations in the pathways detecting these damages. Given the concomitant decreased apoptotic activity, our data support a model in which GR coordinates DNA damage detection and downstream cellular responses, including apoptosis. Thus, our data indicate that GR contributes to the regulation of genome integrity in myeloid cells during muscle regeneration.

At the molecular level, transcriptomic analyses reveal that GR regulates gene networks involved in chromatin organization, DNA repair, and cell cycle progression. However, the comparison with GR cistrome data demonstrates that genes directly bound by GR are not enriched for cell cycle related functions. This apparent discrepancy indicates that GR does not directly control cell cycle genes in myeloid cells, but rather regulates them indirectly. Such indirect regulation is consistent with previous studies showing that GR can modulate transcriptional programs through tethering mechanisms or secondary regulatory cascades, rather than direct DNA binding (^51^ and ref within).

Supporting this model, our interactome analysis uncovers an extensive network of GR associated proteins involved in diverse cellular processes, including cytoskeletal organization, intracellular trafficking, and RNA regulation. Strikingly, GR preferentially associates with HSPA8, a chaperone with ∼85% identity with human HSP70, rather than HSP90 species, indicating that in myeloid cells, GR may predominantly reside in a non-canonical or low affinity conformation^42, 52^. This configuration could therefore limit its classical transcriptional activity and favor non-genomic or protein-protein interaction-based mechanisms. While GR is classically stabilized in an active conformation through HSP90 complexes^42, 52^, its association with HSPA8 has been linked to alternative folding states and intracellular trafficking routes, supporting the idea that GR function in this context may deviate from canonical transcriptional programs. HSPA8 is predominantly localized in the cytoplasm and lysosomal compartments, where it plays a central role in chaperone-mediated autophagy^53, 54^. Through this pathway, it also promotes the degradation of the pro-apoptotic factor BBC3/PUMA under basal conditions, thereby contributing to cellular protection against apoptosis^54^. Moreover, HSPA8 acts as a positive regulator of cell cycle progression by controlling the nuclear accumulation of cyclin D1, a key mediator of the G1 to S phase transition^55, 56^. Consequently, the dissociation of HSPA8 from GR in mutant mice may enhance non-genomic signaling mechanisms to modulate gene expression.

Importantly, our results highlight a strong context- and time-dependent role of GR signaling. Pharmacological activation using dexamethasone reveals that early, but not late, exposure profoundly impairs muscle regeneration. Early glucocorticoid treatment disrupts the transition from monocyte to macrophage, enhances macrophage proliferation, and delays inflammation resolution. These alterations are accompanied by major transcriptional reprogramming affecting cell motility, vascular remodeling, and extracellular matrix organization. While glucocorticoids are widely recognized for their anti-inflammatory properties, their effects on muscle regeneration remain controversial. Previous studies have reported both beneficial and deleterious effects depending on dose, duration, and experimental context^23, 57–60^. Our findings provide a mechanistic framework to reconcile these discrepancies by identifying a critical temporal window during which GR activation is detrimental.

Notably, neutrophils recruitment and cell death appears largely GR-independent at physiological glucocorticoid levels, consistent with previous studies indicating that glucocorticoid responses in neutrophils are primarily related to modulation of trafficking, lifespan and effector functions rather than proliferation^61, 62^. In line with this, neutrophils are terminally differentiated cells with limited proliferative capacity, and several studies have reported rather low or distinct GR activity in these cells.

Together, these findings highlight a cell type-specific role of GR within the myeloid compartment, with a predominant impact on monocyte or macrophage populations^63^. Conversely, upon Dex treatment, our data show an accumulation of neutrophiles 7 dpi. These data resonate with former studies reporting the *in vitro* anti-apoptotic effect of glucocorticoids on human neutrophils, showing that glucocorticoids including Dex inhibit spontaneous neutrophil apoptosis in a concentration-dependent manner (^64^ and ref within).

A particularly unexpected finding is the role of myeloid GR in regulating the crosstalk between macrophages and satellite cells. Early glucocorticoid exposure promotes macrophage proliferation while suppressing satellite cell division, ultimately impairing myogenesis. This observation contrasts with previous studies, which have shown that anti-inflammatory macrophages limit satellite cell proliferation and promote differentiation^8, 26, 65, 66^. In our model, although anti-inflammatory macrophage polarization occurs in both control and mutant conditions, only control macrophages retain the ability to modulate satellite cell proliferation. These findings indicate that acquisition of an anti-inflammatory phenotype based on conventional marker identification is not sufficient to confer functional activity, which instead is highly dependent on GR signaling.

Collectively, our study reveals that GR in myeloid cells does not directly control canonical cell cycle genes, but instead regulates macrophage dynamics through indirect and non-genomic mechanisms involving complex protein interaction networks. Furthermore, it uncovers a GR-dependent intercellular communication axis that is essential for coordinating inflammation resolution and stem cell function. These findings emphasize the importance of timing and environment in glucocorticoid signaling, and highlight the need to reconsider their therapeutic use in regenerative diseases.

## Material and methods

### Mice

Mice were maintained in a temperature- and humidity-controlled animal facility, with a 12-hours light/dark cycle. Standard rodent chow (2800 kcal/kg, Usine d’Alimentation Rationelle, Villemoisson-sur-Orge, France) and water were provided *ad libitum*. Breeding and maintenance of mice were performed according to institutional guidelines. All experiments were done in an accredited animal house, in compliance with French and EU regulations on the use of laboratory animals for research. Intended manipulations were submitted to the Ethical committee (Com’Eth, Strasbourg, France) for approval and to the French Research Ministry (MESR) for ethical evaluation and authorization according to the 2010/63/EU directive under the APAFIS numbers 38761. Dexamethasone (Sigma; product D1756-1G) was dissolved at 20 mg/ml in EtOH, diluted at 0.3 mg/ml in oil, and intraperitoneally administrated at 1 mg/kg. Animals were euthanized *via* cervical dislocation, and tissues were immediately collected, weighed, and frozen in liquid nitrogen or processed for biochemical and histological analyses.

### Generation of GR^LysM-/-^ mice

For myeloid cells specific glucocorticoid receptor (GR) invalidation, GR^L^^2/L^^2^ mice, in which exon 3 (encoding part of the DNA binding domain) was flanked with 2 LoxP sites^31^, were intercrossed with LyzM-Cre mice that express the Cre recombinase selectively in myeloid cells^38^. All mice were on a C57/Bl6J background. Primers used for genotyping are listed in Table S1.

### Histological analyses

For Hematoxylin and eosin staining, tissues were fixed in 10% buffered formalin and embedded in paraffin, as described^67^. Five μm paraffin sections were deparaffinized, rehydrated, and stained in filtered Hematoxylin (Harris, 26041-05) for 3 min, with 1 % Eosin Y (Sigma, 17372-87-1) for 3 min, dehydrated and mounted.

Gomori trichrome staining was performed using a Trichrome Stain Kit according to the manufacturer recommendations with modifications (ScyTek Laboratories). Samples were deparaffinized, rehydrated, and incubated for 1 h in Bouin’s solution preheated to 65 °C under a fume hood. Slides were incubated 10 min in Weigert’s iron hematoxylin stain at room temperature. Excess stain was removed by rinsing in water and acid alcohol solution. Samples were next stained with Trichrome stain solution for 20 min at room temperature, followed by washing in water and acetic acid solution. Slides were dehydrated and mounted. Images were acquired using a NanoZoomer S210 scanner coupled to a Hamamatsu camera.

For immunofluorescent detection and TUNNEL assay, five μm paraffin sections were deparaffinized and rehydrated. Immunofluorescence analysis was performed with antibodies indicated in Table S2, as described^68^. Mouse or rabbit IgGs were used as controls. The quantification of positive cells was performed with QuPath. The various analyses were done with experimenter blinded for the genotype. The TUNEL assay was performed using the *In situ* Cell Death Detection Kit (Roche, 11684795910) according to the manufacturer’s recommendations.

### Fiber cross-sectional area measurements

Muscle cross-sections were stained with Wheat Germ Agglutinin (WGA, Thermofisher W32464) to mark the membrane surrounding each fiber. CSAs were quantified using the FIJI image-processing software. In brief, individual fibers were identified based on the intensity and continuity of the WGA-stained membrane surrounding each fiber by segmentation. Areas were measured after background subtraction, automated thresholding and analyzed with the QuPath software.

### Flow cytometry analysis

Muscle stromal vascular fraction was isolated as described (Ghaibour et al., 2023a). Briefly, hind limb muscles were dissected, minced into small fragments, and enzymatically digested with 1.25 U/mL Dispase II (Stemcell Technologies, Cat: 07913) and 500 µg/mL Collagenase Type I (Thermo Fisher Scientific, 17018029). The resulting cell suspension was filtered through 70 µm cell strainers (Corning Life Sciences) to remove debris. Cells isolated from digested muscle tissue were stained with CD45 Alexa eFluor700 (eBioscience, 56-0451-82, 1:100), CD11b PerCP-Cy5.5 (BioLegend, 101228, 1:100), Ly-6G-GR-1 FITC 488 (BioLegend, 108417, 1:100), Ly-6C PE-CF594 (BD Horizon™, AB_2737749, 1:100), F4/80 APC eFluor 780 (eBioscience™, 47-4801-82, 1:100), and CD206 BUV 785 (BioLegend, 321142, 1:100). Cells were analyzed based on forward scatter (FSC) and side scatter (SSC) parameters to exclude debris and doublets. Data analysis was performed by sequential gating on single, live CD45⁺ leukocytes, followed by identification of myeloid cells as CD11b⁺, neutrophils as CD11b⁺Ly6G⁺, monocytes as CD11b⁺Ly6G⁻Ly6C⁺, and macrophages as CD11b⁺Ly6G-Ly6C-F4/80⁺, with M2-like macrophages defined as CD11b⁺F4/80⁺CD206⁺. Fluorescence intensity was measured using a BD LSR II flow cytometer. Analyses were conducted using FlowJO software, as described ^67^.

### Fluorescence-Activated Cell Sorting (FACS) of Myeloid Cells

After stromal vascular fraction isolation, mononucleated cells were pelleted by centrifugation at 400 × g for 5 min, resuspended in Ham’s F-10 Nutrient Mixture (0.5% Penicillin/Streptomycin solution, and 15% horse serum albumin, without phenol red), and labeled with fluorescence-conjugated antibodies. Cells were stained with CD45 PE (eBioscience, 12-0451-83, 1:100), CD11b PE-Cy7 (eBioscience, 25-0112-82, 1:100). Cells were gated based on FSC and SSC parameters to exclude cellular debris and doublets, and CD45-CD11b double-positive cells were selected. Fluorescence intensity and sorting were conducted using a FACS sorter (BD FACSAria™, BD Biosciences, USA). Sorted satellite cells were subsequently processed for RNA extraction or CUT&RUN analyses.

### RNA extraction and RNAseq analysis

TA muscles were homogenized in TRIzol reagent (Life Technologies, Darmstadt, Germany). RNA was isolated using a standard phenol/chloroform extraction protocol, and quantified by spectrophotometry (Nanodrop, Thermo Fisher). For RNA isolation from FACS-isolated CD11B-positive cells, the RNeasy Plus Micro Kit (Qiagen, 74034) was employed according to the manufacturer’s instructions.

RNA integrity was assessed using a Bioanalyzer. Library preparation was performed at the GenomEast platform from IGBMC. cDNA library was prepared, and sequenced by the standard Illumina protocol (NextSeq 2000, paired-end 50+50 bp for myeloid cells, and single-end 50 bp for TA muscles), following the manufacturer’s instructions. Image analysis and base calling were performed using RTA 2.7.7 and bcl2fastq 2.17.1.14. Adapter dimer reads were removed using DimerRemover (-a AGATCGGAAGAGCACACGTCTGAACTCCAGTCAC). FastQC 0.11.2 (http://www.bioinformatics.babraham.ac.uk/projects/fastqc/) was used to evaluate the quality of the sequencing. Reads were aligned to the mouse mm10 and mm39 genome for TA muscles and myeloid cells, respectively, using STAR v2.7.10b^69^, retaining only uniquely mapped reads for downstream analyses. Only uniquely aligned reads were retained for further analyses. Quantification of gene expression was performed using HTSeq 0.11.0^70^. For comparison among datasets, the transcripts with more than 50 raw reads were considered.

Differentially expressed genes (DEGs) were identified using the Bioconductor libraries DESeq2^71^ with a p < 0.05. Identified DEGs were subjected to functional analysis using STRING 12.0^72^ and to pathway analysis using WebGestalt^73^ with the Over-Representation Analysis (ORA) method, applying a false discovery rate (FDR) < 0.05 as the significance threshold.

Heatmaps were generated by centering and normalizing expression values using Cluster 3.0 ^74^, with subsequent visualization Morpheus (https://software.broadinstitute.org/morpheus/). Genes were clustered using K-Mean with gene tree construction, Pearson correlation, and average linkage as clustering parameters ^75^.

### CUT&RUN analysis

CUT&RUN analysis has been performed on FACS-isolated myeloid cells as described^76^. Briefly, FACS-isolated satellite cells were centrifuged at 500 × g for 10 min at 4°C, and the resulting cell pellet was incubated with nuclear extraction buffer (20 mM HEPES-KOH pH = 7.9, 10 mM KCl, 0.5 mM Spermidine, 0.1% Triton X-100, 20% Glycerol and protease inhibitor cocktail) for 20 min on ice. Isolated nuclei were then washed and incubated with Concanavalin A-coated beads (BioMag Plus Concanavalin A, Cat: 86057-3) for 10 min at 4°C. Following additional washes, the nuclei-bead complexes were incubated overnight at 4°C with gentle agitation in the presence of 5 µg of primary antibodies targeting GR (C-terminal, IGBMC, #3249), H3K27ac (active motif 39133) and H3K4me3 (Abcam, ab1012-100), or a rabbit IgG (Santa Cruz, sc2357) negative control. Nuclei-bead complexes were precipitated and incubated with protein A-micrococcal nuclease (pA-MN, IGBMC) for 1 h at 4°C under constant agitation. Enzymatic cleavage was initiated by adding 3 µL of 100 mM CaCl₂, followed by 30 min of incubation on ice. The reaction was terminated at 37°C for 20 min using stop buffer (200 mM NaCl, 20 mM EDTA, 4 mM EGTA, 50 µg/mL RNase A, 40 µg/mL glycogen, 25 pg/mL yeast spike-in DNA). DNA was extracted from immunoprecipitated samples for sequencing.

Libraries were sequenced with an Illumina Hiseq 2000 as paired-end 50 bp reads, and mapped to the mm10 reference genome using Bowtie 1.3.1^77^. Uniquely mapped reads were retained for further analysis. Reads overlapping with ENCODE blacklisted region V2 were removed using Bedtools ^78^, and remaining reads into two groups: fragment size <120 bp (without nucleosome, in general for transcription factors) and fragment size >150 bp (with nucleosomes, normally for histone marks). Bigwig files were generated using bamCoverage (deeptools 3.3.0: bamCoverage --normalizeUsing RPKM --binSize 20), and raw bedgraph files with genomeCoverageBed (bedtools v2.26.0). SEACR 1.3 algorithm (stringent option) was used for the peak calling with a threshold of 0.002. The genome-wide intensity profiles were visualized using the IGV genome browser (http://software.broadinstitute.org/software/igv/)^79^ or Figeno^80^. HOMER was used to annotate peaks and for motif searches^81^. *De novo* identified motifs were referred to as follow: R = purine (G or A); Y = pyrimidine (T or C). Genomic features (promoter/TSS, 5’ UTR, exon, intron, 3’ UTR, TTS and intergenic regions) were defined and calculated using Refseq and HOMER according to the distance to the nearest TSS.

Venn diagrams were generated with Venny 2.1.0 (https://bioinfogp.cnb.csic.es/tools/venny/) or InteractiVenn^82^. Pathway analysis was performed with WebGestalt using the Over-Representation Analysis (ORA) method^73^.

Clustering analyses were performed using seqMINER ^83^ and deeptools (-computeMatrix -skip0; - plotHeatmap). Genomic intersections and sequence extractions were carried out using bedtools. GetFastaBed was used to extract FASTA sequences from BED files. Intersect Interval was applied to identify overlapping genomic locations. When not specified, all bioinformatics analyses were performed using default parameters, ensuring robust and reproducible results.

Pearson correlation analysis with deeptools to determine the similarity between the samples^84^. Use the command line multiBamSummary bins --bamfiles file1.bam file2.bam -o results.npz, followed by plotCorrelation -in results.npz --corMethod pearson --skipZeros --plotTitle “Pearson Correlation of Read Counts” --whatToPlot heatmap --colorMap RdYlBu --plotNumbers -o heatmap_PearsonCorr_readCounts.png --outFileCorMatrix PearsonCorr_readCounts.tab.

### scRNAseq analyses

PRJNA577458 was used for scRNA-seq analysis of mouse muscles 2, 3.5, 5 and 10 dpi, using metacluster annotations, as described^85^. Bioinformatics analyses were assessed with Seurat v5.0.1. Data normalization was performed with -NormalizeData, and scaled with -ScaleData with default features. Myeloid cells were clustered according to *Itgam*/*Cd11b* expression. Loupe file was generated with the -create_loupe_from_seurat function from the loupeR package v1.0.2. Loupe Browser 8 was used for colocalization analyses.

### Caspase activity

The caspase activity was determined on three controls and three mutants’ batches of about 5000 FACS-purified neutrophils, monocytes or macrophages each, using the Caspase-Glo 3/7 assay (Promega, USA).

### Immunoprecipitation and Liquid Chromatography Tandem Mass Spectrometry (LC-MS/MS)

FACS-isolated myeloid cells were harvested in PBS and centrifugated at 300 × *g* for 5 min. Pellets were lysed for 30 min under agitation in lysis buffer [Tris (pH 7.5) 50 mM, NaCl 150 mM, glycerol 5%, NP-40 1%, PIC 1× in H20], as described^86^. Immunoprecipitation was performed using Slurry Dynabeads G coupled to 5 μg of anti-GR antibody or a rabbit IgG for 1 h at 4°C under agitation. On ice, 100 μg of proteins were added on antibody-coupled beads and agitated at 4◦C overnight. After three washes in lysis buffer, bound proteins were eluted in 1x Laemmli and 0.1 M DTT and boiled. Isolated proteins were subjected to an overnight trypsin digestion at 37°C. Peptides were separated on a C18 column using an Ultimate 3000 nano-RLSC (Thermo Fisher Scientific) coupled to an LTQ-Orbitrap Elite mass spectrometer (Thermo Fisher Scientific) for peptide identification. Proteins were identified with Proteome Discoverer 2.4 software (Thermo Fisher Scientific, PD2.4 from Mus Musculus proteome database (Swissprot), and proteins were quantified with a minimum of two unique peptides based on the XIC (sum of the Extracted Ion Chromatogram). Partners of GR were considered if enriched (p<0.05, |log fold-change| > 0.5) in IP-GR condition compared to the protein listed in the IgG sample to exclude non-specific bindings. For visual representation and Gene Ontology terms analysis, the online tool STRING.db^72^ was used.

## Statistical analyses

Data processing and analysis were conducted using DESEq2, Microsoft Excel and GraphPad Prism 9. In all graphical representations, error bars indicate the standard error mean (SEM). Comparisons between groups were performed using appropriate statistical tests, selected based on the number of groups, data distribution (normality), variance homogeneity, and multiple comparisons. The specific statistical tests applied are indicated in the figure legends. Statistical significance was defined as follows: ns, p > 0.05 (non-significant); *, p < 0.05; **, p < 0.01; and ***, p < 0.001.

## Resource availability

Custom macro for CSA measurements quantification is available on GitLab (CSA: <u>the BIOP cellpose</u> <u>extension guide</u>). All sequencing datasets generated in this study are deposited in the Gene Expression Omnibus (GEO) with the accession number GSE311683 for CUT&RUN analyses, GSE311684 and GSE311685 for RNA-seq experiments.

## Acknowledgments

We thank Dr. R. Ricci (IGBMC) for generously providing the LysM-Cre mouse line. We are grateful to Anastasia Bannwarth, Brayann Weis, and Régis Lutzing for their valuable technical support, and Tao Ye, Matthieu Jung and Céline Keime for excellent bioinformatics assistance. We also thank members of flow cytometry IGBMC facilities Claudine Ebel, Muriel Philipps, Cécile Macquin and Virginie Sutter for their technical support. We acknowledge the support of the IGBMC core facilities, including the animal facility, microscopy and mass spectrometry platforms, cell culture service, flow cytometry, and histopathology services, as well as the Mouse Clinical Institute (ICS, Illkirch-Graffenstaden, France) and GenomEast facility, a member of the ‘France Génomique’ consortium (ANR-10-INBS-0009). We are grateful to Jan Tuckerman, Bénédicte Chazaud, Rémi Mounier and Elodie Monsellier for helpful discussions.

This work was supported by the Interdisciplinary Thematic Institute IMCBio as part of the ITI 2021-2028 program of the University of Strasbourg, the Centre National pour la Recherche Scientifique (CNRS), Institut national de la santé et de la recherche médicale (Inserm), from IDex Unistra (ANR-10-IDEX-0002), SFRI-STRAT’US (ANR 20-SFRI-0012) and EUR IMCBio (ANR-17-EURE-0023) projects, under the framework of the French Investments for the Future Programme. Additional support was provided by Inserm, CNRS, Unistra, IGBMC, Agence Nationale de la Recherche and Deutsche Forschungsgemeinschaft (ANR-10-BLAN-1108, ANR-DFG CORTICOSAT, ANR-22-CE92-0020-01). J.G.R. received funding from the Programme CDFA-07-22 from the Université franco-allemande and Ministère de l’Enseignement Supérieur de la Recherche et de l’Innovation, E.C. by MESRI, and R.S. by IMCBio.

## Author contributions

D.D., D.M., and S.S.C. formulated the initial hypothesis. S.S.C., J.G.R., E.Ca., E.Co., I.C., and Q.C. carried out functional, molecular, and histological experiments. D.D., V.G., and S.S.C. executed bioinformatics analyses. S.S.C. and E.G. took responsibility of imaging analyses. R.S. and B.M. took responsibility for mass-spectrometry analysis. S.S.C., D.M., and D.D. took primary responsibility for data analysis and writing the manuscript.

## Declaration of generative AI and AI-assisted technologies

None declared.

