## Supplementary Figures 1 to 3 and Tables 1 to 3 for "Glucocorticoid Receptor Signaling in Myeloid Cells Orchestrates Inflammation Resolution and Muscle Repair"

This file includes Supplementary Figures 1-3 and Supplementary Tables 1-3.

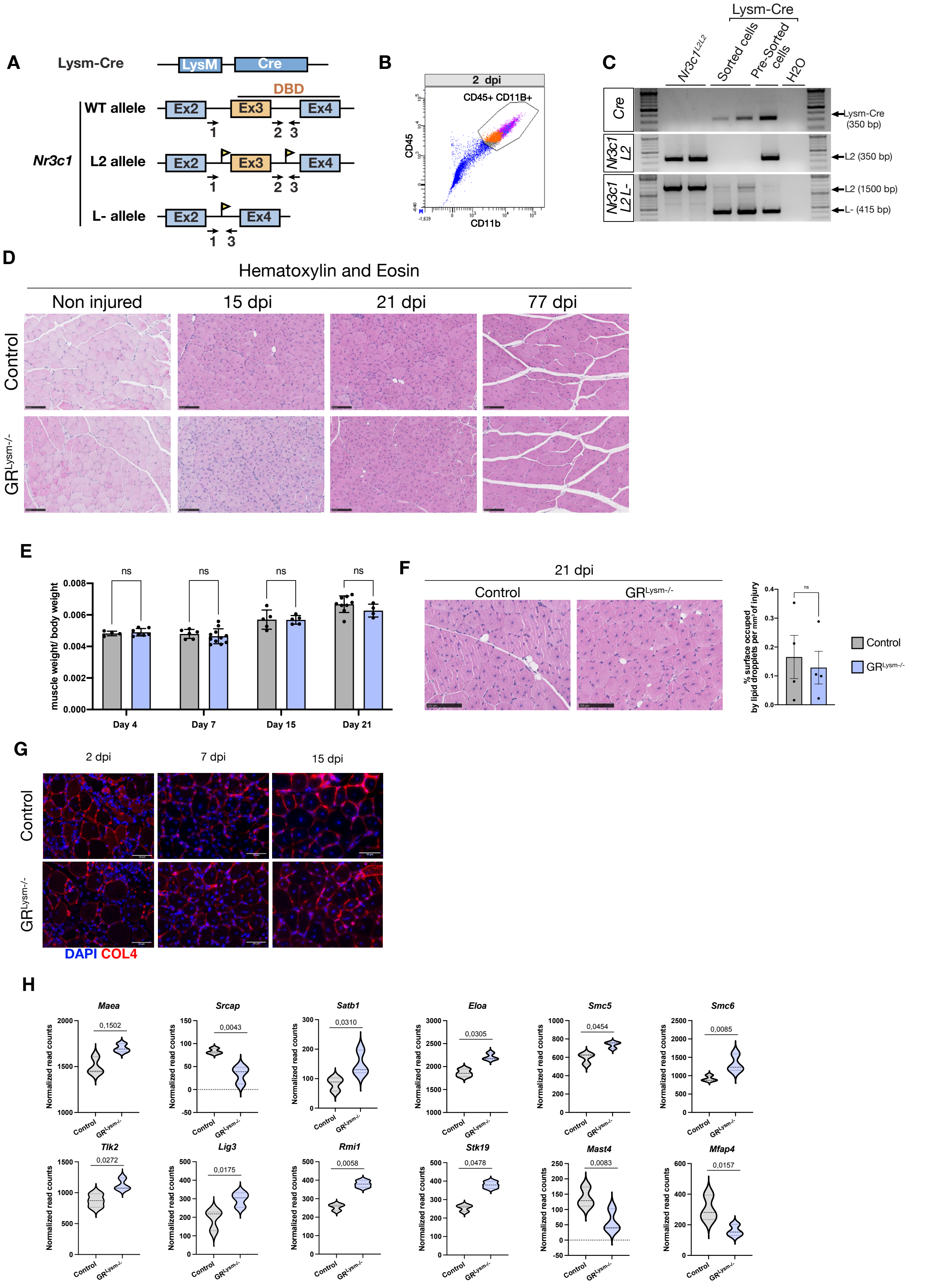

**Figure S1: Efficient invalidation of GR in myeloid cells**

(A) Schematic representation of *LysM*-Cre, wild-type (WT), floxed (L2) and Cre-mediated exon 3 deleted (L-) *Gr* alleles. LoxP sites are shown by arrowheads. Primers used for allele characterization are depicted by arrows and sequences are in material and method section.

(B) Representative flow cytometry density plot showing the gating strategy used to isolate $CD45^+/CD11b^+$ myeloid cells from injured TA muscles.

(C) Representative electrophoresis of PCR-amplified genomic DNA from control and $GR^{Lysm-/-}$ mice. Distinct amplicon sizes correspond to Cre (350 bp), GR L2 (1500 bp or 350 bp), and the recombined GR L− allele (415 bp). The absence of the GR L− band in $NR3C1^{L2/2}$ control mice confirms their genotype, whereas its presence in both FACS-sorted $CD45^+CD11b^+$ and presorted cell populations demonstrates successful recombination in the mutant.

(D) Representative hematoxylin and eosin staining of TA muscles of control and $GR^{Lysm-/-}$ mice under non-injured condition, and at 15-, 21- and 77-days post-injury (dpi). Scale bars, 100 µm.

(E) Muscle weight normalized to total body weight in injured mice at 4-, 7-, 15- and 21-days post-injury (dpi). Muscle mass was measured at the indicated time points following injury and normalized to each animal's total body weight. Data are presented as mean ± SEM. Statistical test used was Student's t-test; ns, non-significant.

(F) Representative hematoxylin and eosin staining of 21 days-injured TA muscles from control and $GR^{Lysm-/-}$ mice showing lipid droplet accumulation within the injury site. Quantification represents the surface area of lipid droplets relative to the total injury surface. Data are presented as mean ± SEM. Statistical test used was Student's t-test; ns, non-significant.

(G) Representative immunofluorescent detection of COL4 (red) of TA of control and $GR^{Lysm-/-}$ mice at indicated time points of muscle regeneration. Scale bar, 15 µm.

(H) Violin plots showing normalized read counts of selected genes that were differentially expressed in bulk RNA-sequencing of FACS-sorted $CD11b^+$ cells isolated from TA muscles of control and $GR^{Lysm-/-}$ mice 7 dpi.

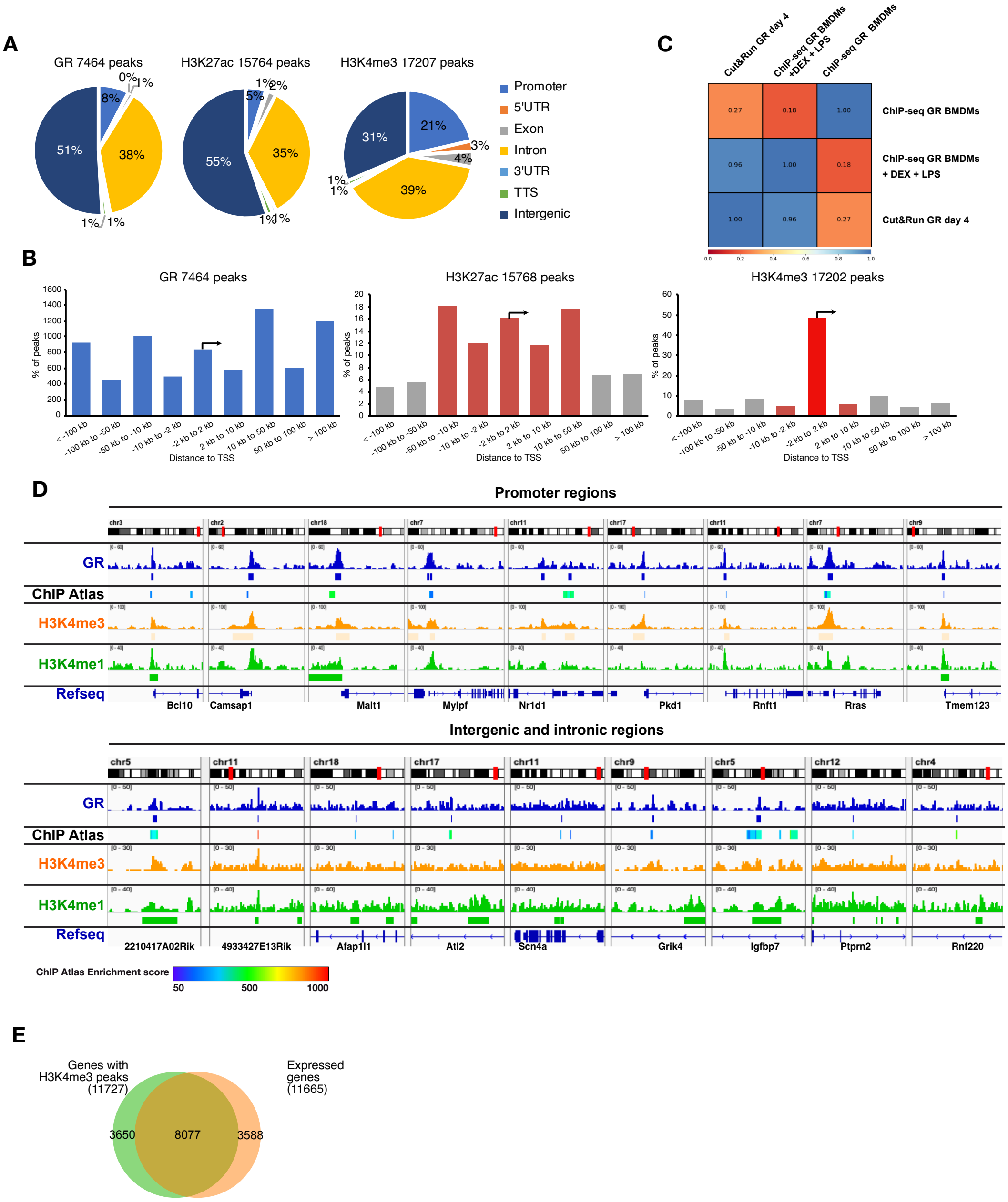


**Figure S2: Integrative chromatin profiling of GR in myeloid cells isolated from TA muscle 4 dpi**

(A) Pie chart depicting the genomic location of H3K4me3, H3K27ac and GR in macrophages.

(B) Distribution of H3K4me3, H3K27ac and GR peaks relative to their distance to the nearest TSS.

(C) Pearson correlation of GR in macrophages CUT&RUN analysis with public ChIP-seq datasets.

(D) Localization of GR, H3K4me3 and H3K27ac at representative promoter and intergenic regions on the chromatin of macrophages by CUT&RUN. Enrichment score showing the levels of confidence of published datasets is depicted as a heatmap.

(E) Overlap of the genes bound by H3K4me3 in macrophages expressed genes in myeloid cells.

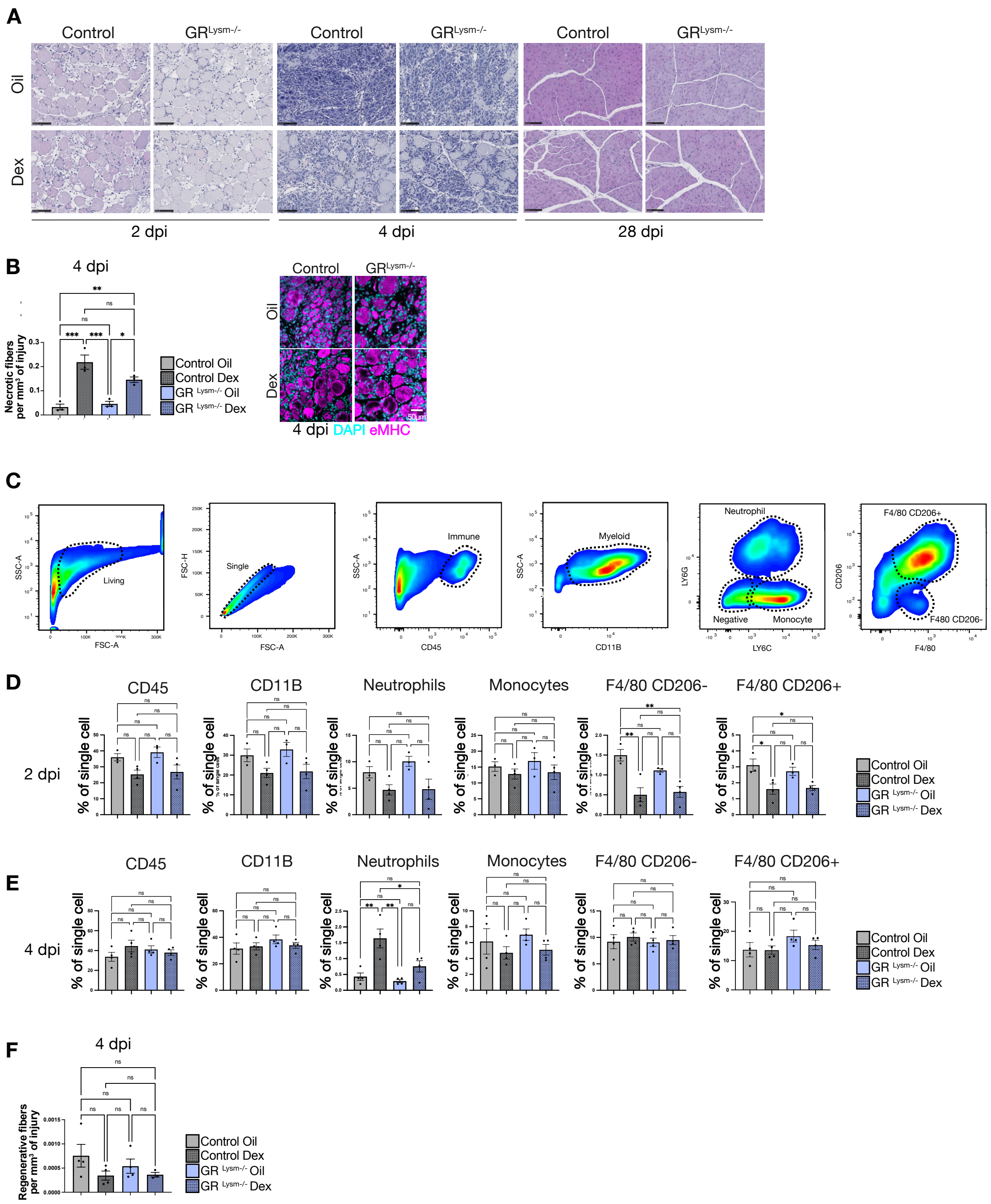


**Figure S3: Dexamethasone treatment during the pro-inflammatory phase impairs early muscle regeneration**

(A) Representative hematoxylin and eosin staining of tibialis muscles (TA) of dexamethasone- (Dex) or vehicle-treated (Oil) control and GR$^{Lysm-/-}$ mice 2, 4 and 28 days after injury. Scale bar, 100 µm.

(B) Representative immunofluorescent detection of eMHC (magenta) in tibialis muscles of Dex- or Oil-treated control and GR$^{Lysm-/-}$ mice 4 days after injury. Quantification represents the percentage of necrotic fibers relative to the injury surface. Statistical test used is Two-way ANOVA with Tukey *post-hoc* correction. The various comparisons were non-significant. Scale bar, 50 µm.

(C) Representative flow cytometry gating strategy used to identify myeloid populations in injured TA muscles. $CD45^+$ cells were first selected as immune cells, and $CD11b^+$ cells as myeloid cells. Neutrophils were defined as $Ly6G^+CD11b^+$ cells, monocytes as $Ly6C^+/Ly6G^-/CD11b^+$ cells, and macrophages as $Ly6G^-/Ly6C^-/CD11b^+/F4\text{-}80^+$ cells further subdivided into $CD206^-$ (M1-like) and $CD206^+$ (M2-like) subsets.

(D) Percentage of myeloid cells in 2-days-injured TA of Dex- or Oil-treated control and $GR^{Lysm-/}$. Data are presented as mean ± SEM. Statistical test used is Two-way ANOVA with Tukey *post-hoc* correction; ns, non-significant; * = $p < 0.05$; ** = $p < 0.01$; *** = $p < 0.001$.

(E) Percentage of myeloid cells in 4-days-injured TA of Dex- or Oil-treated control and $GR^{Lysm-/}$. Data are presented as mean ± SEM. Statistical test used is Two-way ANOVA with Tukey *post-hoc* correction; ns, non-significant; ** = $p < 0.01$; *** = $p < 0.001$.

(F) Quantification represents the percentage of regenerative fibers relative to the injury surface in 4-days-injured TA of Dex- or Oil-treated control and $GR^{Lysm-/-}$mice. Data are presented as mean ± SEM. Statistical test used is Two-way ANOVA with Tukey *post-hoc* correction; ns, non-significant; ** = $p < 0.01$; *** = $p < 0.001$

**Supplementary Tables**

**Supplementary Table 1: Primers for mouse genotyping**

| Gene | Forward | Reverse |
|---|---|---|
| *LysMCre* | 5’ TTCCCGCAGAACCTGAAGATGTTCG 3’ | 5’ GGGTGTTATAAGCAATCCCCAGAAATGC 3’ |
| *GRL2/WT* | 5’ AGATCATTTGCCTAGCAGGCATGAG 3’ | 5' GTCAACACATGATCACCTTGCAGTC 3' |
| *GRL-* | 5’ CCAGAGAACTAATTGGCICTIGCAC 3’ | 5' GTCAACACATGATCACCTTGCAGTC 3' |

**Supplementary Table 2: Antibody list**

| Antibody / Dye | Reference |
|---|---|
| LY6C/G | Abcam AB2557 |
| IBA1 | Abcam AB5076 |
| F4/80 | Cell signaling 70076 |
| KI67 | R&D systems 14-5698-82 |
| gH2AX | Sigma-Aldrich clone JBW301 |
| WGA | W32464 thermofisher |
| MYOG | Abcam AB1835 |
| PAX7 | Thermo Fisher PA1-117, |
| ACTN1 | Abcam ab9465 |
| COL4 | Sigma AB769 |
| CD163 | Abcam EPR19518, ab182422 |
| CD45 | Abcam AB10558 |
| eMHC | Dshb F1.652 |
| GR | Home made |

**Supplementary Table 3: List of GR partners identified by immunoprecipitation followed by mass-spectrometry**

See Table S3.xlsx
